# An HMA-like integrated domain in the wheat tandem kinase WTK4 recognises an RNase-like pathogen effector

**DOI:** 10.1101/2025.08.26.672365

**Authors:** Zoe Bernasconi, Ursin Stirnemann, Yang Xu, Aline G. Herger, Renjie Chen, Victoria Widrig, Matthias Heuberger, Mariantonietta Lettieri, Marion C. Müller, Brande B. H. Wulff, Thomas Wicker, Beat Keller, Javier Sánchez-Martín

## Abstract

Proteins with a tandem kinase structure have recently emerged as new players in race-specific resistance in cereal crops. However, the molecular understanding of these novel immune receptors’ resistance mechanisms is limited by the lack of knowledge about the pathogen effectors that they recognise. In this work, we identify AvrWTK4, the wheat powdery mildew RNase-like effector recognised by the wheat tandem kinase immune receptor WTK4, through a combination of bi-parental genetic mapping and mutagenesis. We demonstrate that mutations in the *AvrWTK4* gene or a reduction of its expression lead to virulence on *WTK4*. Transfection of *AvrWTK4* specifically induced cell death in *WTK4*-containing *Aegilops tauschii* protoplasts. The avirulent AvrWTK4 variant interacts more strongly than the virulent variant with the N-terminal heavy metal-associated (HMA)-like domain of WTK4. These findings further highlight that integrated domains in tandem kinase proteins serve as decoys for pathogen effectors, which could be leveraged to design novel recognition specificities.

## Introduction

Cereal crops such as wheat and barley are essential components of the human diet, yet their production is threatened by pathogens that reduce agricultural yields (Savary et al. 2019). Plants have an innate immune system composed of different layers; the first layer is activated by the recognition of conserved pathogen-associated molecular patterns (PAMPs) via membrane-bound receptors, leading to PAMP-triggered immunity (PTI) (DeFalco and Zipfel 2021). Adapted pathogens have evolved mechanisms to bypass PTI by secreting effectors, which repress the immune response and facilitate infection. However, plants have a second layer of defence, activated via recognition of effectors, then called avirulence (Avr) factors, by intracellular resistance (R) proteins, resulting in effector-triggered immunity (ETI) (Dodds and Rathjen 2010; Ngou, Ding, and Jones 2022). ETI is race-specific and frequently results in hypersensitive response (HR) and cell death.

Most known *R* genes encode nucleotide-binding leucine-rich repeat receptors (NLRs) (Kourelis and van der Hoorn 2018). In addition, kinase fusion proteins have recently been identified as a novel class of R proteins involved in ETI in wheat and barley (Klymiuk et al. 2021; Liu, Hou, and Chen 2024). These non-canonical *R* genes encode proteins with a domain organisation made of a kinase domain fused to either another kinase domain, forming what is known as tandem kinase proteins (TKPs; some examples being *Rpg1*, *Yr15 =* WTK1*, Sr60 =* WTK2*, Pm24/RWT4 =* WTK3*, WTK4* and *Sr62* = WTK5), and/or to domains of diverse nature (for example, *Pm4*) and have important roles in conferring race-specific resistance to various biotrophic fungal diseases, such as different rust species, powdery mildews and blast (Liu, Hou, and Chen 2024; Chen, Gajendiran, and Wulff 2024).

Some NLR immune receptors carry integrated domains (IDs), which act as “effector traps” by mimicking host targets of pathogen effectors (Kroj et al. 2016; Cesari 2018). By resembling these native targets, IDs enable direct interaction with effectors that manipulate host metabolism or immunity (Marchal et al. 2022; Cesari 2018; Bailey et al. 2018). Well-characterised examples include the rice NLRs RGA5 and Pikp-1, both of which feature heavy metal-associated (HMA)-like IDs that bind the rice blast effectors AVR-Pia/AVR1-CO39 and AVR-Pik, respectively (Kanzaki et al. 2012; Cesari et al. 2013; Maqbool et al. 2015). Notably, HMA-like IDs have recently emerged as promising targets for engineering immune receptors with expanded effector recognition (Zdrzałek et al. 2024). Interestingly, recent work has shown that over half of plant TKPs also carry IDs, suggesting they may function similarly to ID-carrying NLRs, by acting as decoys for pathogen effectors (Reveguk et al. 2025).

In powdery mildew of wheat (*Blumeria graminis* f. sp. *tritici*; *Bgt*) and barley (*B. hordei*; *Bh*), numerous studies have focused on identifying and characterising *Avr* effectors recognised by NLR immune receptors. Notable examples include the *AvrPm2* and *AvrPm3* effectors in *Bgt* (Bourras et al. 2015; Bourras, Kunz, Xue, Praz, Muller, et al. 2019; Praz et al. 2017), and the *Avra* effectors in *Bh* (X. Lu et al. 2016), all exhibiting a typical RNase-like structure. Genetic approaches such as GWAS and bi-parental mapping have been instrumental in identifying *Avr* genes in both pathosystems (Praz et al. 2017; Bourras et al. 2015; Kunz et al. 2023; Müller et al. 2022). Recently, UV mutagenesis led to the discovery of *AvrPm4*, the first *Bgt* effector recognised by a kinase fusion protein (Bernasconi et al. 2025), as well as the avirulence regulator *Bgt-646*, whose disruption results in virulence on both the NLR Pm3a and the TKP WTK4, suggesting a shared regulatory pathway (Bernasconi et al. 2024). *Avr* effectors from both *Bgt* and *Bh* typically trigger cell death when co-expressed with their corresponding *R* gene in *Nicotiana benthamiana* or wheat/barley protoplasts (Saur, Bauer, Lu, et al. 2019; Bourras et al. 2016; Bernasconi et al. 2025). In *Bh*, Avra effectors directly interact with their corresponding Mla receptors (Saur, Bauer, Kracher, et al. 2019). In *Bgt*, AvrPm3^b2/c2^ was recently found to associate with the LRR domain of Pm3b (Isaksson et al. 2025), while a predicted wheat zinc finger protein has been identified as a mediator of the Pm2-AvrPm2 interaction, forming a tripartite complex essential for immune activation (Manser et al. 2024).

Although kinase fusion proteins have emerged as a novel class of immune receptors mediating race-specific resistance, their underlying resistance mechanisms remain poorly understood (Sánchez-Martín and Keller 2021; Athiyannan et al. 2022). Among the few characterised examples, AvrPWT4, the *Magnaporthe oryzae* effector recognised by the TKP RWT4 (Inoue et al. 2017; Arora et al. 2023), was recently shown to interact with RWT4 via a newly identified ID, formed by a partial duplication of the RWT4’s kinase domain, KDup (Sung et al. 2025), suggesting a possible role for IDs as effector decoys in TKPs. Furthermore, we recently demonstrated that AvrPm4 directly interacts with, and is phosphorylated by, the kinase fusion protein Pm4 (Bernasconi et al. 2025). Recent studies indicate that TKP-mediated resistance may depend on additional host components. For instance, both SR62 and RWT4 require a helper NLR to activate immune responses in the presence of avirulent AvrSr62 and AvrPWT4 variants, respectively (Chen et al. 2025; P. Lu et al. 2025). Despite these advances, our understanding of how pathogen effectors evade recognition by TKP immune receptors is limited. Identifying these Avr effectors and investigating how they evolve to evade TKP-mediated recognition are key steps towards evaluating resistance durability and guiding the development of more effective and sustainable resistance strategies (Tosa and Chuma 2014; Lovelace et al. 2023).

In this study, we combined bi-parental mapping and UV mutagenesis to identify *AvrWTK4*, encoding an RNase-like effector of wheat powdery mildew recognised by the TKP WTK4. We showed that point mutations and modulation of *AvrWTK4* expression, mediated by the ankyrin-repeat protein Bgt-646, are the main mechanisms to overcome *WTK4*-mediated resistance. We identified an HMA-like ID at the N-terminus of WTK4 and showed that it directly interacts with AvrWTK4. Modifications of the binding residues, either in the HMA-like ID or in AvrWTK4, led to reduced interaction. Our findings reveal a TKP which mediates resistance through an integrated decoy mechanism. This may represent an evolutionarily conserved strategy among tandem kinase immune receptors and potentially offers a framework for engineering these immune receptors in cereals to gain expanded or novel recognition specificity.

## Results

### Bi-parental mapping in wheat powdery mildew identifies a single *AvrWTK4* locus on chromosome 5

To understand at the molecular level the resistance function of the previously cloned *R* gene *WTK4* (Gaurav et al. 2022), we aimed at identifying its corresponding avirulence (*Avr*) gene by combining bi-parental genetic mapping and AvrXpose, a UV mutagenesis-based approach for *Avr* gene identification (Bernasconi et al. 2024).

For bi-parental mapping, we used a pre-existing F1 mapping population derived from a cross between a *Bgt* isolate, CHE_96224, and a *B. g. triticale* isolate, THUN-12 (Müller et al. 2019). We observed that THUN-12 was virulent on 21 *Ae*. *tauschii* lines containing *WTK4*, on which CHE_96224 was avirulent (**Supp. Table S1**; (Gaurav et al. 2022)). This indicated that THUN-12 did not harbour *AvrWTK4*, making the CHE_96224 x THUN-12 mapping population (Müller et al. 2019) suitable to map the *AvrWTK4* locus. Three *Ae. tauschii* lines TOWWC0028, TOWWC0040, and TOWWC0111 were randomly selected from the 21 lines mentioned above, and 83, 70, and 70 randomly chosen progeny of the *Bgt* mapping population were used for phenotyping on the three lines, respectively (**Supp. Table S2**). For all three lines, a single locus for avirulence on *WTK4* was identified on chromosome 5 (**Fig. 1A**). The locus interval varied between the three different *Ae. tauschii* lines used; however, the region of best association was overlapping (18,756,380 Mb to 19,163,249 Mb; hereafter, *AvrWTK4* locus) and contained the best associated SNP for all the three lines (**Fig. 1B**). In the *AvrWTK4* locus, there were 25 annotated genes, among which three were polymorphic between THUN-12 and CHE_96224. Two of these three genes, *Bgt-55150* and *BgtE-5764,* were annotated as effector genes, encoding proteins with a predicted signal peptide (**Fig. 1B**; **Supp. Table S3**). On the protein level, Bgt-55150_THUN-12_ differs from Bgt-55150_96224_ by three amino acids (T30N, T33P, and E35Q), while BgtE-5764_THUN-12_ differs from BgtE-5764_96224_ by one amino acid (V255A). Bgt-55150 is 119 amino acids in length and highly expressed, whereas BgtE-5764 is 257 amino acids in length and is expressed at a low level at the haustorial stage in CHE_96224 (Praz et al. 2018). Based on these characteristics, we considered *Bgt-55150* the most likely *AvrWTK4* candidate gene. Interestingly, the previously cloned gene *AvrPm3^b2/c2^* (*BgtE-20002*) is also in the *AvrWTK4* locus; however, it is not polymorphic between THUN-12 and CHE_96224, which are both avirulent on *Pm3b* (Bourras, Kunz, Xue, Praz, Müller, et al. 2019; Müller et al. 2021). Therefore, it was not considered for further investigation.

**Fig. 1.**
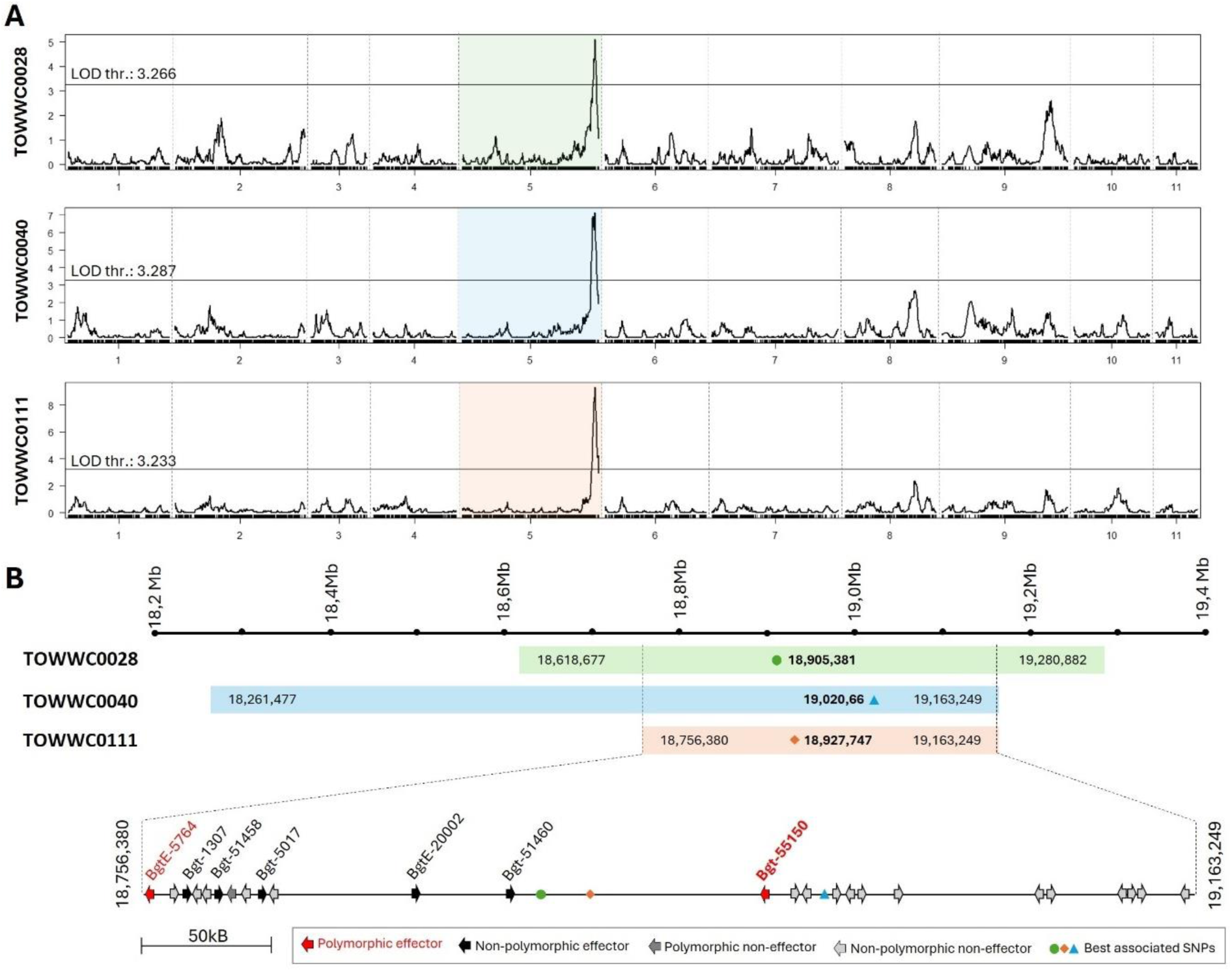
Identification of the *AvrWTK4* locus using a bi-parental mapping population. (**A**) QTL mapping in a bi-parental mapping population (CHE_96224 x THUN-12 (Müller et al. 2019)) on three different *Ae. tauschii* lines containing *WTK4* identified a single locus for avirulence on *WTK4* on chromosome 5 (coloured). The LOD (logarithm of the odds) threshold, shown as a black line, was calculated using 1,000 permutations. (**B**) The locus interval varied between the three different *Ae. tauschii* lines (TOWWC0028, TOWWC0040, and TOWWC0111); however, their regions of best association overlapped. The best-associated SNPs for the *Ae. tauschii* lines are located within the common region of association, between 18,756,380 and 19,163,249 Mb in chromosome 5 of the CHE_96224 genome (*AvrWTK4* locus). These SNPs are represented as a dot (TOWWC0028), a triangle (TOWWC0040), and a square (TOWWC0111) on the plot. The LOD scores at the position of the best-associated SNP were 5.086 (TOWWC0028), 7.135 (TOWWC0040), and 9.291 (TOWWC0111). Among the 25 annotated genes within the *AvrWTK4* locus, seven are annotated as effectors (Müller et al. 2019), and two of them, *BgtE-5764* and *Bgt-55150*, are polymorphic between CHE_96224 and THUN-12 (indicated with a red arrow; see also **Supp. Table S2**).

### *WTK4* gain of virulence mutants have mutations in the *Avr* candidate *Bgt-55150*

To confirm the *Avr* candidate found through bi-parental mapping as *AvrWTK4*, we employed the UV mutagenesis-based approach AvrXpose (Bernasconi et al. 2024) to generate *Bgt* gain-of-virulence mutants on *WTK4*. We generated six *Bgt* mutants virulent on *WTK4* (hereafter, *AvrWTK4* mutants), which were single-spore isolated and used for phenotyping on *WTK4*-containing *Ae. tauschii* lines (TOWWC0087, TOWWC0112, and TOWWC0154), as well as on a set of *Pm* differential lines to determine their virulence specificity. The *AvrWTK4* mutants were found to be virulent on the three tested *Ae. tauschii* lines containing *WTK4*, but not on any of the other tested *Pm* differential lines (**Fig. 2A**), except for mutant AvrWTK4_5, which was also virulent on *Pm3f* (**Supp. Fig. S1**). Whole genome sequencing and comparison to the non-mutated parental isolate CHE_96224 revealed that three *AvrWTK4* mutants (namely, AvrWTK4_3, AvrWTK4_4, and AvrWTK4_6) had a single nucleotide polymorphism (SNP) in the *AvrWTK4* candidate mapped above, *Bgt-55150*. Surprisingly, AvrWTK4_3 and AvrWTK4_6 shared the same mutation (T30N; **Supp. Table S4-S5**). These mutations were the result of independent events, as the mutants were isolated in separate mutagenesis experiments and showed a different set of additional mutations across the whole genome (**Supp. Tables S4-S5**). Of note, the T30N mutation present in mutants AvrWTK4_3 and AvrWTK4_6 also occurs in the virulent variant of Bgt-55150 (Bgt-55150_THUN-12_), suggesting that this amino acid position is particularly relevant for specific recognition by WTK4. Mutant AvrWTK4_4 had a SNP in the predicted intron sequence of *Bgt-55150* (43 bp from the 5’ splice site and 13 bp from the 3’ splice site). We used primers binding on the two exons to PCR amplify a part of *Bgt-55150* from cDNA of isolates CHE_96224 and AvrWTK4_4 and observed that the size of the PCR product was larger in AvrWTK4_4 (**Fig. 2B**), suggesting that the intron of *Bgt-55150* is retained in AvrWTK4_4. This was then confirmed by sequencing the PCR product. The loss of splicing leads to a premature stop codon in the translated protein. None of the *AvrWTK4* mutants had mutations in the coding sequence of *BgtE-5764,* the other polymorphic effector in the *AvrWTK4* locus found with bi-parental mapping (**Supp. Table S4-S5**). Mutant AvrWTK4_2 had a SNP in the putative 3’ regulatory region of *BgtE-5764*, 531 base pairs (bp) downstream of the stop codon; however, based on transcriptomic data of CHE_96224 (Praz et al. 2018) showing that the transcript level is very low at position 480 bp and absent at 670 bp after the stop codon of BgtE-5764, we concluded that the SNP has no effect on gene expression and that the causal mutation is likely affecting a different gene.

**Fig. 2.**
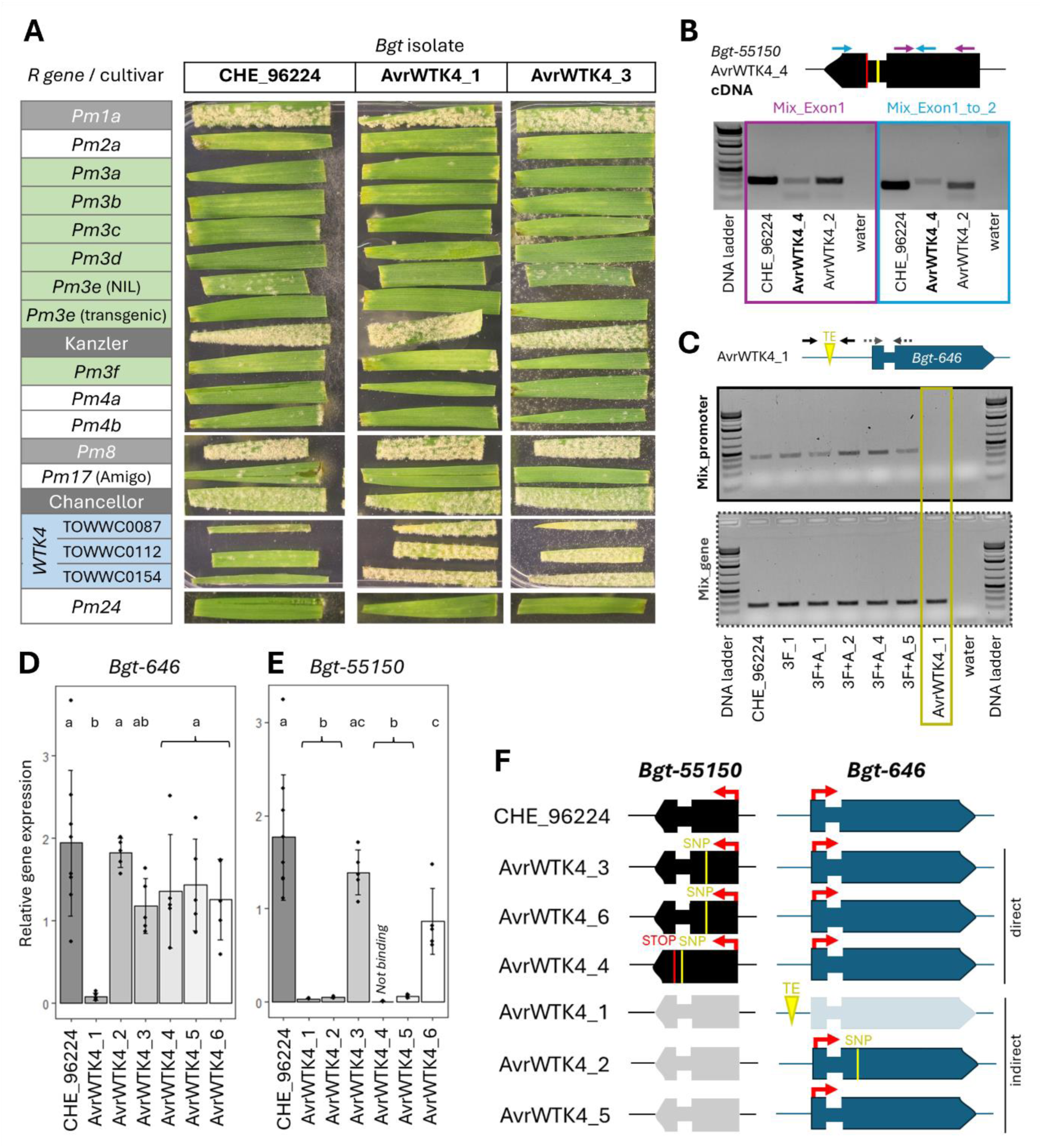
The *AvrWTK4* mutants exhibit direct or indirect gain-of-virulence mechanisms, all affecting *Bgt-55150*. (**A**) Wheat lines containing different *R* genes were infected with CHE_96224 and the *AvrWTK4* mutants. Cultivars susceptible to CHE_96224 (the *Pm1a* NIL, the sister line of the transgenic *Pm3e* line (here called *Se#2*), the *Pm8* NIL and Chancellor) are highlighted in grey. NILs containing *Pm3* alleles are depicted in green, and the three *Ae. tauschii* lines containing *WTK4* are in blue. The remaining *Pm* lines are shown in white. The phenotype of two representative mutants, AvrWTK4_1 and AvrWTK4_3, is shown (the phenotype of the other *AvrWTK4* mutants can be found in **Supp. Fig. S1**). (**B**) PCR amplification at the cDNA level of *Bgt-55150* shows an increased size of the central gene part (Mix_Exon1_to_2) in AvrWTK4_4 compared with CHE_96224 and AvrWTK4_2 (220 base pairs instead of the 165 expected), indicating that the intron is retained. (**C**) PCR amplification of the *Bgt-646* gene (grey dashed arrows) and promoter region (black arrows) in CHE_96224, five selected 3F and 3F+A mutants (virulent on *Pm3a* and *WTK4*; see previous study (Bernasconi et al. 2024)) and mutant AvrWTK4_1. No amplification was observed for AvrWTK4_1 at the site of the TE insertion, indicating that the TE insertion is too large for amplification under the extension time used. The drawing of *Bgt-646* is not to scale. Expression analysis of (**D**) *Bgt-646* and (**E**) *Bgt-55150* in the *AvrWTK4* mutants. One of the primers for *Bgt-55150* detection spans over the two exons (**Supp. Table S6**). Thus, for mutant AvrWTK4_4, where splicing is not occurring due to an intron mutation, the primer is not binding. The observed lack of relative gene expression confirms the intron retention. Significance and p values (significance threshold of p < 0.05) were calculated using a two-sided analysis of variance (ANOVA) followed by a Tukey’s HSD test (**Supp. Table S7**). Tests with the same letter are not significantly different. (**F**) Schematic representation of the genes *Bgt-55150* and *Bgt-646* in the *AvrWTK4* mutants. Direct or indirect changes in sequence or expression of *Bgt-55150* are indicated. Brighter boxes indicate genes with low or no expression, while expressed genes are darker boxes marked with a red arrow. SNPs and TE insertions are in yellow. Gene representations are not to scale.

### Mutations in the effector regulatory gene *Bgt-646* result in a reduction of *AvrWTK4* expression and virulence on *WTK4*

As mutants AvrWTK4_1, AvrWTK4_2, and AvrWTK4_5 did not have mutations in *Bgt-55150*, we reasoned that additional components involved in avirulence on *WTK4* must exist. We investigated whether these three mutants had any commonly mutated genes and discovered that two of them had mutations affecting the gene *Bgt-646*, encoding an ankyrin-repeat-containing protein. Mutant AvrWTK4_1 had an insertion of a transposable element 580 bp upstream of the start of the coding sequence (CDS) of *Bgt-646*, causing a strong reduction in *Bgt-646* expression (**Fig. 2C-D**) and mutant AvrWTK4_2 had a SNP in the CDS of *Bgt-646,* causing an amino acid change (R93K; **Supp. Tables S4-S5**).

Interestingly, in a previous study, we observed an additional gain of virulence on *WTK4* in *Bgt* mutants with gain of virulence on *Pm3a*, an NLR immune receptor (Bernasconi et al. 2024). Three of these previously described *Pm3a/WTK4* virulent mutants also had mutations in *Bgt-646*, which was then shown to regulate the expression of several effector genes in *Bgt*, including *Bgt-55150* (Bernasconi et al. 2024). To confirm whether this regulatory role extends to the *AvrWTK4* mutants, we performed an expression study and observed a significantly reduced expression of *Bgt-55150* in both *AvrWTK4* mutants with mutations in *Bgt-646*, suggesting that *Bgt-646* may affect *Bgt-55150* expression (**Fig. 2E**). Finally, mutant AvrWTK4_5 did not have mutations in *Bgt-646,* or any other commonly mutated gene, but *Bgt-55150* expression was also repressed (**Fig. 2E**). Thus, we hypothesise that additional, yet unidentified, mutations in AvrWTK4_5 reduce the expression of *Bgt-55150*, causing the gain of virulence. These findings further support our previous study (Bernasconi et al. 2024), indicating that mutations in *Bgt-646* result in lower expression of *Bgt-55150* and cause a loss of avirulence on *WTK4*.

In conclusion, all *WTK4*-virulent *Bgt* mutants have either direct (via SNPs) or indirect (via reduced expression) modifications of *Bgt-55150* (**Fig. 2F**). These data, combined with the bi-parental mapping approach, strongly indicate that *Bgt-55150* is *AvrWTK4*.

### *WTK4*-mediated resistance in *Ae. tauschii* induces a hypersensitive response

NLRs that confer resistance to biotrophic pathogens such as *Bgt* in most cases induce HR, leading to cell death and ultimately stopping the pathogen from spreading. To investigate whether the immune response mediated by *WTK4* also results in HR, we performed a microscopic evaluation of resistance upon infection. The avirulent isolate CHE_96224 was used to infect two *WTK4-*containing *Ae. tauschii* lines (TOWWC0028 and TOWWC0077), and the fungal infection was observed at 24, 48, and 96 hpi. In both *Ae. tauschii* lines, at least 70% of the spores attempting to penetrate were blocked at a similar rate by both HR and papilla formation at all three timepoints (**Fig. 3A**-**B**; **Supp. Fig. S2**). These data provide evidence that the *WTK4*-induced immune response results in both HR and papilla formation, very similar to other *Pm* resistance genes such as *Pm4*, encoding a kinase fusion protein (Sánchez-Martín et al. 2021).

**Fig. 3.**
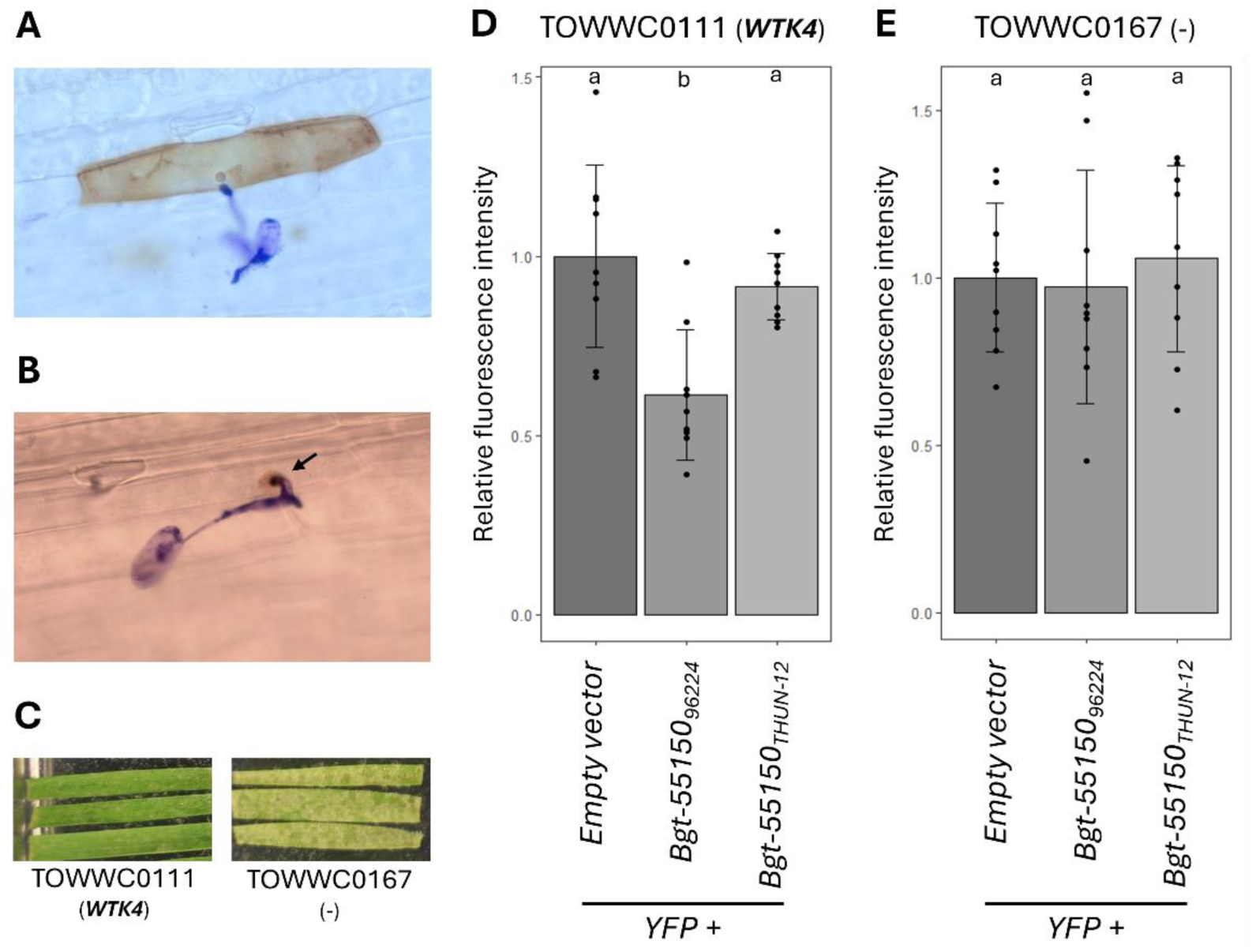
*WTK4*-mediated cell death in *Ae. tauschii* depends on *Bgt-55150_96224_*. At the microscopic level, inoculation of a *WTK4*-containing *Ae. tauschii* line with CHE_96224 results in **(A)** hypersensitive response (brown colouring, visualised using DAB staining), and (**B**) papilla formation (round structure indicated by a black arrow), with cell-to-cell variation within the same line. The quantification of the different types of defence response is described in **Supp. Fig. S2**. (**C**) The *Ae. tauschii* line TOWWC0111 (containing *WTK4*) is resistant, while TOWWC0167 is susceptible 7 days post inoculation with the *Bgt* isolate CHE_96224. (**D**) Protoplasts derived from TOWWC0111 and (**E**) TOWWC0167 were co-transfected with 3 pmol of a vector encoding YFP and 3 pmol of either an empty vector, *Bgt-55150_96224_* (avirulent allele) or *Bgt-55150_THUN-12_* (virulent allele). YFP fluorescence was measured 18-20 hpi for two technical and three biological replicates per treatment. The experiment was repeated three times, for a total of 9 biological replicates. Fluorescence values were normalised to the average fluorescence of protoplasts co-transfected with the YFP vector and the empty vector for each experiment. Significance and p values (significance threshold of p < 0.05) were calculated with a two-sided analysis of variance (ANOVA) followed by a Tukey’s HSD test (**Supp. Table S6**). Boxes with the same letter are not significantly different.

To functionally validate *Bgt-55150_96224_* as *AvrWTK4*, we used a cell death assay relying on co-transfection of *Ae. tauschii* protoplasts, a method commonly used to functionally validate *Avrs* that trigger HR upon recognition (Saur, Bauer, Kracher, et al. 2019; Arndell et al. 2024; Chen et al. 2025). The *Bgt-55150* gene sequences were codon-optimised for plant expression and cloned into expression vectors without signal peptide. We selected a resistant (TOWWC0111, containing *WTK4*) and a susceptible (TOWWC0167, lacking *WTK4*) *Ae. tauschii* line for this experiment (**Fig. 3C**). Co-transfection of *Bgt-55150_96224_* with a YFP marker in the *Ae. tauschii* line TOWWC0111 significantly reduced YFP fluorescence intensity by 33% compared to *Bgt-55150_THUN-12_* co-transfected with the YFP marker (**Fig. 3D**). In TOWWC0167, no significant difference in relative YFP fluorescence intensity was observed between protoplasts transfected with either an empty vector, *Bgt-55150_96224_* or *Bgt-55150_THUN-12_* (**Fig. 3E**), indicating that *Bgt-55150_96224_* induces cell death specifically in the presence of *WTK4*. From these results we conclude that *Bgt-55150_96224_* is *AvrWTK4*.

To check if all necessary components for the cell death response are conserved among other plant species, we performed cell death assays in *N. benthamiana* and in wheat protoplasts. Intriguingly, co-expression of *WTK4* with *Bgt-55150_96224_* in *N. benthamiana* did not lead to cell death (**Supp. Fig. S3A**). Additionally, protoplasts of the wheat cultivar Fielder (which lacks *WTK4*) co-transfected with *WTK4* and *Bgt-55150_96224_* did not show significantly reduced relative YFP fluorescence intensity compared to protoplasts co-transfected with *WTK4* and an empty vector or *Bgt-55150_THUN-12_*, indicating that no cell death occurred (**Supp. Fig. S3B**). These findings indicate that at least one genetic component unique to *Ae. tauschii* and absent in *N. benthamiana* or the wheat cultivar Fielder is needed for inducing HR upon recognition of *Bgt-55150_96224_*.

### WTK4 interacts with AvrWTK4 through its HMA-like integrated domain

To assess whether WTK4 and Bgt-55150_96224_ (hereafter, AvrWTK4_96224_) interact at the protein level, we first modelled their 3D protein structures using AlphaFold2 (Mirdita et al. 2022). AvrWTK4_96224_ is predicted to exhibit the typical RNase-like fold of all *Bgt Avrs* identified to date (**Fig. 4B**) (Bourras et al. 2016; Bourras, Kunz, Xue, Praz, Müller, et al. 2019), while WTK4, in addition to the pseudokinase (K1) and kinase (K2) domains, also has a predicted N-terminal heavy metal-associated (HMA)-like domain, which has not been previously described (Gaurav et al. 2022; **Fig. 4A**-**B**; amino acid residues 1-68; hereafter HMA_WTK4_). HMA_WTK4_ shows relatively low amino acid identity to HMA-like integrated domains (IDs) of known NLR proteins (i.e. RGA5 and Pikp-1; 30.8 and 32.4% amino acid sequence identity, respectively; **Supp. Fig. S4**; Cesari et al. 2013; Maqbool et al. 2015), and the closest homolog in *Arabidopsis thaliana* is the HIPP protein HIPP39 (51.5% amino acid sequence identity between the two HMA-like domains). Nevertheless, using the 3D model of HMA_WTK4_ to assess structural homology using TM-align (Zhang 2005), we observed that HMA_WTK4_ showed very high (> 82%) structural homology to the HMA-like domains of RGA5, Pikp-1 and HIPP39 (**Table 1**). To extend this analysis, we investigated whether other TKPs also have HMA-like IDs that may have gone unnoticed in previous studies. We found that RPG1, the first TKP identified in barley (Brueggeman et al. 2002), also has an HMA-like ID at its N-terminus (52.9% sequence identity with HMA_WTK4_ at the protein level, and 93% at the structural level; **Supp. Fig. S4**, **Table 1**). None of the other TKPs examined showed any additional domain with structural similarity to HMA_WTK4_ (**Supp. Fig. S5**).

**Fig. 4.**
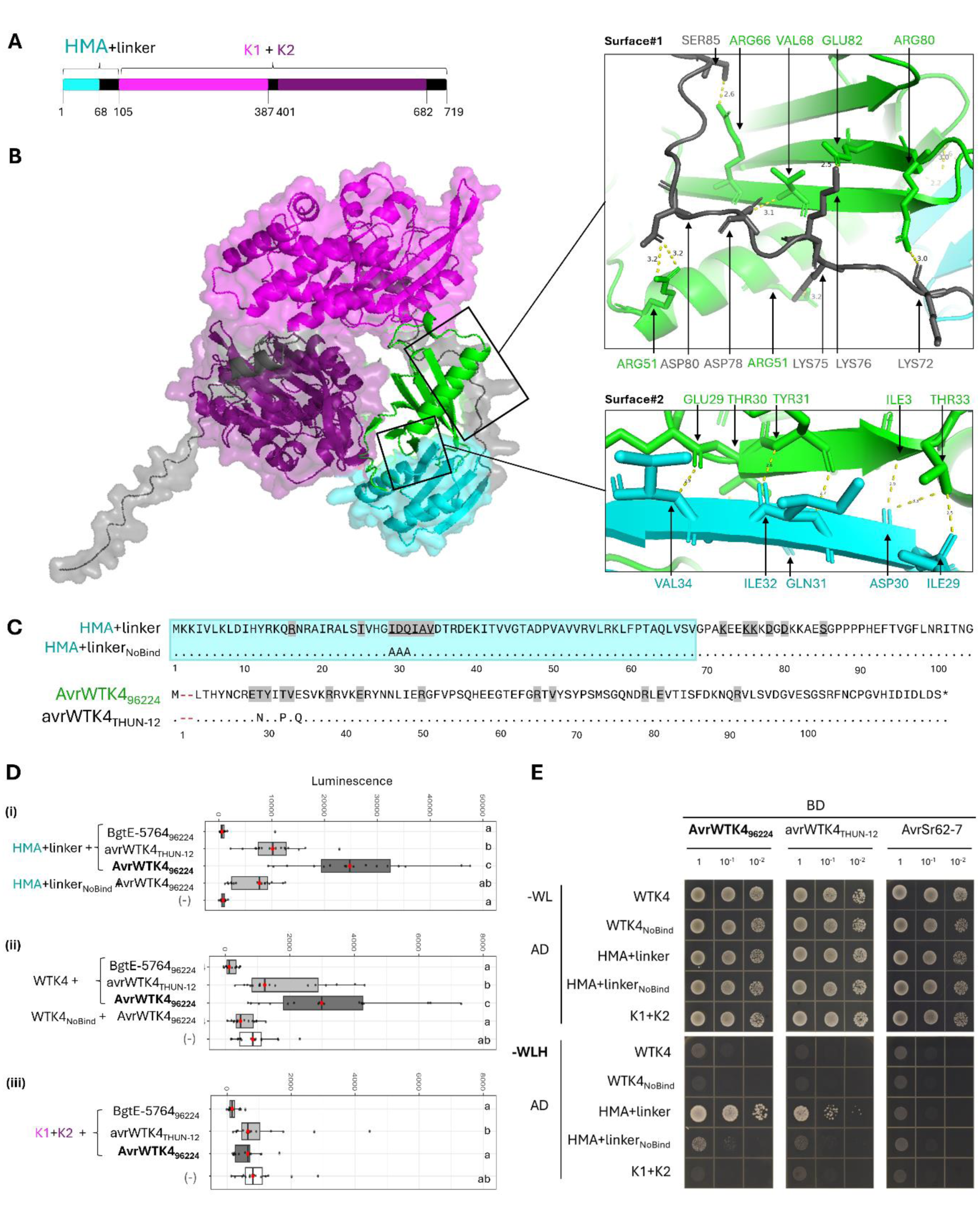
WTK4 interacts with AvrWTK4 through its HMA-like integrated domain. (**A**) Schematic view of WTK4 protein domains predicted by ProSite, with their amino acid positions indicated: HMA-like domain (cyan), pseudokinase (K1, light purple), kinase (K2, deep purple), and unstructured sequence (black). (**B**) AvrWTK4_96224_ (green), featuring an RNase-like fold (based on AlphaFold3 modelling) is predicted to interact with WTK4 through two main interaction surfaces: Surface#1, involving the linker sequence, and Surface#2, involving the HMA-like ID. Residues involved in each interaction, along with distances (in ångström), are shown in the corresponding images. (**C**) Protein sequences of the HMA-like ID with the linker sequence (HMA+linker), a modified version with three amino acid changes and reduced binding affinity to AvrWTK4_96224_ (HMA+linker_NoBind_), and two variants of Bgt-55150 (AvrWTK4_96224_ and avrWTK4_THUN-12_) are shown. Signal peptides are omitted (two red dashes). Residues involved in interactions are highlighted in grey and underscored. (**D**) The interaction of AvrWTK4_96224_, avrWTK4_THUN-12_, and BgtE-5764_96224_ was tested with i) HMA_WTK4_ fused to the linker sequence (HMA+linker), ii) WTK4 full-length, and iii) the two kinase domains of WTK4 (K1+K2). avrWTK4_THUN-12_ and BgtE-5764_96224_ interacted less than AvrWTK4_96224_ with both the full-length WTK4 and the HMA_WTK4_+linker domain. The modified constructs WTK4_NoBind_ and HMA+linker_NoBind_ showed significantly reduced luminescence relative to their non-mutated versions. The negative control for all experiments is AvrWTK4_96224_ with the NLR protein PM17 (Müller et al. 2022). Significance and p values (significance threshold of p < 0.05) were calculated with a two-sided analysis of variance (ANOVA) followed by a Tukey’s HSD test (**Supp. Table S7**). Boxes with the same letter are not significantly different. (**E**) A yeast-2-hybrid (Y2H) assay shows significantly stronger interaction of AvrWTK4_96224_ with HMA+linker compared to avrWTK4_THUN-12_. AvrWTK4 variants were fused to the GAL4 binding domain (BD) and WTK4 variants to the GAL4 activation domain (AD). Yeast suspensions at an OD_600_ = 1.0 and two serial dilutions of 1:10 and 1:100 were spotted on synthetic medium lacking tryptophan and leucine (-WL, growth control) and on synthetic medium lacking tryptophan, leucine, and histidine (-WLH, interaction selection). Immunoblot of each protein is shown in **Supp. Fig. S8**. The negative control, AvrSr62-7, did not interact with any of the constructs tested.

**Table 1.** Structure of selected proteins with HMA-like domains and their similarity to HMA_WTK4._

| Gene | Species of origin | Resistance | Structure | TM score - HMA<sub>WTK4</sub><sup>1</sup> |
| --- | --- | --- | --- | --- |
| *WTK4* | Wheat | Wheat powdery mildew | TKP | 1 |
| *RPG1* | Barley | Barley stem rust | TKP | 0.93065 |
| *RGA5* | Rice | Rice blast | NLR | 0.83728 |
| *Pikp-1* |  |  | NLR | 0.82739 |
| *HIPP39* | *Arabidopsis* | none | HIPP | 0.94011 |

<sup>1</sup> Proteins with a TM score higher than 0.6 (HMA<sub>WTK4</sub> against target sequence) are considered structural homologues. A TM score of 1 indicates 100% structural similarity of the two compared 3D structures.

As IDs in NLRs have previously been described to be involved in direct interactions with fungal effectors (Cesari 2018), we hypothesised that HMA_WTK4_ physically interacts with AvrWTK4_96224_. To test this, we used AlphaFold Multimer to model the complex. The predicted ipTM value, indicative of model confidence, was 0.73 for the AvrWTK4_96224_-WTK4 (full length) and 0.86 for the AvrWTK4_96224_-HMA_WTK4_ interaction, suggesting a stable association. The model depicted two main interaction surfaces on WTK4: one involving HMA_WTK4_ and one the linker sequence between HMA_WTK4_ and the pseudokinase domain, K1 (**Fig. 4B**). Notably, the T30N substitution found in the virulent isolate THUN-12 and in mutants AvrWTK4_3 and AvrWTK4_6 (**Fig. 1C**) is located within the predicted surface #2 (**Fig. 4 B-C**). In fact, modelling the interaction between WTK4 and the virulent Avr variant, Bgt-55150_THUN-12_ (hereafter, avrWTK4_THUN-12_) yielded a much lower ipTM value of 0.17, indicating a loss of interaction (**Supp. Fig. S6**). This supports the hypothesis that only the avirulent AvrWTK4_96224_ variant stably interacts with HMA_WTK4_, and that the mutations present in the virulent isolate likely disrupt this interaction.

We next validated these interactions *in vivo* using a split-luciferase assay in *N. benthamiana.* Our results showed that AvrWTK4_96224_ specifically interacts with HMA_WTK4_ fused to the linker sequence (HMA+linker; **Fig. 4D i**). AvrWTK4_96224_ also interacted with full-length WTK4, but to a lesser extent, likely due to its lower protein levels (**Fig. 4D ii**, **Supp. Fig. S7**). In contrast, the virulent avrWTK4_THUN-12_ variant showed significantly reduced interaction with both constructs, indicating that amino acid residues 30, 33, and 35 in *AvrWTK4* are critical for binding. Protein levels of both effectors were comparable (**Supp. Fig. S7**). Both AvrWTK4_96224_ and avrWTK4_THUN-12_ showed very weak interaction with the kinase domains of WTK4 in the absence of HMA_WTK4_ (K1+K2), with luminescence levels similar to the negative control (**Fig. 4D iii**). Changing three residues (I29, D30, Q31) predicted to form the interaction surface #2 in HMA_WTK4_ to three alanine residues (**Fig. 4C**) led to a significant drop in luminescence for both the full-length WTK4 (WTK4_NoBind_) and the HMA+linker (HMA+linker_NoBind_) constructs, confirming the functional relevance of this interface (**Fig. 4D i-ii**). No interaction was detected between WTK4 constructs and BgtE-5764_96224_, the other polymorphic effector from the *AvrWTK4* locus (**Fig. 4D**).

These results were corroborated by yeast two-hybrid assays: strong interaction was observed between AvrWTK4_96224_ and HMA+linker, while the virulent avrWTK4_THUN-12_ variant showed considerably weaker interaction (-WLH; **Fig. 4E**). The HMA+linker_NoBind_ protein, with modified Avr binding interface, showed no interaction with AvrWTK4_96224_ (**Fig. 4E**). No interaction was detected between any WTK4-derived protein and the negative control AvrSr62-7, an unrelated effector recognised by the TKP SR62^TK^ (Chen et al. 2025). BgtE-5764, the *Bgt* effector used as negative control in the split luciferase assay, was autoactive in yeast, thus, it could not be used in the yeast two-hybrid assay (**Supp. Fig. S8**).

These findings demonstrate that the HMA-like domain of WTK4 mediates direct recognition of AvrWTK4. Alongside insights from the gain-of-virulence *Bgt* mutants, we conclude that virulent *Bgt* isolates evade recognition either through transcriptional downregulation of *AvrWTK4*, or via point mutations that specifically affect its HMA-binding interface, reducing direct interaction with the immune receptor. Overall, our results reveal a mechanistic basis for the recognition of effectors by tandem kinase proteins via integrated decoy domains, similar to ID-containing NLRs. This could lay the groundwork for further exploration of TKP integrated domains as platforms for engineering immune receptors with broader resistance spectra.

## Discussion

We identified *AvrWTK4*, the first powdery mildew effector mediating avirulence on a tandem kinase protein (TKP), using two complementary approaches: bi-parental mapping and mutagenesis. *AvrWTK4* encodes an RNase-like effector, and the pathogen can evade WTK4-mediated recognition by mutating key amino acid residues or by downregulating its gene expression. Importantly, we show that recognition depends on an integrated decoy domain in WTK4 that mediates direct interaction with AvrWTK4.

It is noteworthy that two independent *AvrWTK4* mutants have the exact same mutation in *Bgt-55150*, resulting in the T30N substitution at the protein level, a mutation that is also present in the naturally virulent isolate THUN-12. Structural modelling and interaction assays confirmed that this mutation strongly reduces interaction with the HMA domain of WTK4 (**Fig. 4**; **Supp. Fig. S6**). Other studies, mostly done with NLR immune receptors, have shown that single point mutations in pathogen effectors allow escape from recognition. For instance, the globally widespread AvrPm8_F43Y variant effectively evades recognition by Pm8 (Kunz et al. 2023).

In addition to amino acid changes, downregulation of expression of *AvrWTK4* represents a second gain-of-virulence mechanism. Our data suggest that mutations resulting in downregulation occur at similar frequencies as amino acid changes: in three out of six mutants, the *AvrWTK4* sequence was unchanged, but its expression was significantly reduced (**Fig. 2**). In two of these cases (*Bgt* mutants AvrWTK4_1 and AvrWTK4_2), we identified mutations in *Bgt-646*, encoding an ankyrin-repeat protein previously described as a regulator of expression of several effector genes (Bernasconi et al. 2024). As described in this earlier work, modifications of *Bgt-646* conferred virulence on both *Pm3a* and *WTK4*, suggesting a shared regulatory pathway (Bernasconi et al. 2024). Here, we confirm that *Bgt-646* indirectly determines *WTK4* virulence by controlling *AvrWTK4* expression. In various pathosystems, including *Bgt*, additional genetic components to Avrs, such as suppressors of Avr recognition, have been shown to influence R-Avr interactions (Petit-Houdenot and Fudal 2017; Bourras et al. 2016; Bernasconi et al. 2025; Bourras et al. 2015). However, in contrast to suppressors that block recognition, *Bgt-646* appears to act as a positive regulator of effector expression. Transcriptional regulators of virulence genes have been described in other fungal species (John et al. 2021). Whether disruption of regulators of effector gene expressions is a widespread gain-of-virulence mechanism in *Bgt* remains to be elucidated, but it appears to be common at least in the cases of recognition by Pm3a, an NLR (Bernasconi et al. 2024), and WTK4, a TKP (this study). Further experiments are needed to elucidate the molecular mechanism by which *Bgt-646* regulates effector gene expression, and how many genes are controlled by *Bgt-646*.

Intriguingly, in the case of the mutant AvrWTK4_5, the mutation causing the virulence phenotype remains unknown, as no mutations were found in either *AvrWTK4* or *Bgt-646* (**Supp. Tables S4-S5**). This is consistent with our previous study, in which we could not find the phenotype-causing mutation for three out of seven *Pm3a/WTK4* virulent mutants (Bernasconi et al. 2024). This suggests a more complex regulatory network controlling *AvrWTK4* expression and points to additional, yet unidentified, components involved in modulating avirulence on *WTK4*.

Infections in *Ae. tauschii* lines with a *WTK4*-avirulent *Bgt* isolate triggered a hypersensitive response (HR) (**Fig. 3, Supp. Fig. S2**), and *WTK4*-expressing *Ae. tauschii* protoplasts showed partial but significant cell death upon transfection with *AvrWTK4_96224_* (**Fig. 4**). This is similar to observations in *WTK3*-containing wheat landraces, where mild cell death occurred after infection with an avirulent *Bgt* isolate (P. Lu et al. 2020). By contrast, *WTK4*-*AvrWTK4_96224_* co-expression failed to induce HR in *N. benthamiana* or wheat (cv. Fielder) protoplasts, suggesting that *WTK4* requires additional *Ae. tauschii*-specific components. This resembles the Pm4-AvrPm4 interaction, which triggers HR in wheat but not in *N. benthamiana* (Bernasconi et al. 2025).

In NLR-mediated immunity, sensor NLRs with integrated decoy domains often require helper NLRs to fully activate defence responses, as described for RGA5/RGA4 in rice or RPS4/RRS1 in *Arabidopsis* (Baggs, Dagdas, and Krasileva 2017; Jubic et al. 2019; Cesari et al. 2013; Narusaka et al. 2009). Likewise, TKPs such as SR62^TK^ and RWT4 were shown to require a helper NLR to mediate resistance to stem rust and wheat blast, respectively (Chen et al. 2025; P. Lu et al. 2025). Similarly, in the case of the tandem kinase gene *Yr15*, different introgression lines showed varying resistance phenotypes, indicating that *Yr15* is part of a larger resistance complex involving additional genetic components (Klymiuk et al. 2018, 2020). Given that WTK4 has an ID that directly binds to the AvrWTK4 effector, we hypothesise that it might act as a sensor, for which the putative helper NLR is not yet known, and whose presence would trigger HR in *WTK4-AvrWTK4* co-expression assays. Together, these findings support a broader model in which TKPs act as sensors that depend on helper proteins (possibly NLRs) for full immune activation. To find the additional component required for *WTK4* function, interaction screens, such as yeast-two-hybrid or immunoprecipitation followed by mass spectrometry (IP-MS), or extensive mutant analyses in search of second-site mutations may be performed.

The discovery of an HMA-like ID in WTK4 represents a significant advance in understanding the mechanistic basis of resistance conferred by TKPs, a novel class of immune receptors. A recent study found that over half of TKPs across the plant kingdom carry IDs (Reveguk et al. 2025). Moreover, the presence of IDs in at least four characterised resistance proteins with a TKP structure (WTK4, RPG1, Lr9, and RWT4; this study; Sung et al., 2025; Wang et al., 2023), suggests a common, functional role in pathogen recognition. However, the specific contributions of these domains remain largely unexplored. In the case of RWT4, KDup is directly involved in the interaction with the *M. oryzae* effector AvrPWT4 (Sung et al. 2025). Lr9 has an additional vWA domain, but to our knowledge, this ID has not yet been functionally characterised (Wang et al. 2023). Our findings reinforce the emerging view that IDs within TKPs may play central roles in effector recognition.

Several effectors known to interact with integrated HMA-like domains of NLRs, such as AVR-Pia, AVR1-CO39, and AVR-Pik, belong to the MAX (*Magnaporthe* Avrs and ToxB-like) effector family, a structurally conserved group common in *M. oryzae* (Seong and Krasileva 2021; Cesari et al. 2013; Maqbool et al. 2015). *WTK4* confers resistance against wheat powdery mildew, a pathogen which lacks MAX-like effectors in its repertoire (Seong and Krasileva 2023, 2021). Its corresponding effector, AvrWTK4, belongs to the RNase-like family and has no sequence or structural similarity with MAX effectors (Seong and Krasileva 2023; Bourras et al. 2016). This shows that effectors from different pathogen species have convergently evolved to target and bind HMA domains from their hosts. This, in turn, suggests that HMA(-like) domains integrated in immune receptors might be capable of evolving diverse effector-binding specificities, possibly through minimal sequence changes. It remains to be explored whether different effector types bind overlapping or distinct surfaces on HMA domains. Such knowledge could potentially be used to design synthetic HMA modules capable of recognising a broader range of Avr effectors, even from unrelated pathogens, such as *Bgt* and *M. oryzae*. Here, we predicted and then experimentally confirmed a specific interaction between HMA_WTK4_ and the avirulent AvrWTK4 variant, but not with the virulent variant, illustrating the potential of AlphaFold for *in silico* analysis of recognition specificity (**Supp. Fig. S6**).

So far, engineering efforts have shown that altering a few residues in the HMA domain of RGA5 to mimic those of Pikp-1 enables RGA5 to recognise AVR-Pik, thereby broadening its recognition specificity (Cesari et al. 2022). Similarly, replacing the HMA domain of Pikp-1 with that of a natural host target, HIPP43, conferred broad-spectrum resistance (Zdrzałek et al. 2024). In parallel, nanobody-based IDs have been introduced into NLRs to expand recognition potential, offering a promising approach for engineering immune receptors (Kourelis et al. 2023). These studies show that the identification and study of IDs in plant immune receptors are of high relevance, illustrating their potential as platforms for designing immune receptors with broadened or redirected specificity.

In conclusion, we demonstrated that the identification of novel Avr effectors is instrumental to understand resistance mechanisms of nonconventional immune receptors. Identifying and characterising *AvrWTK4* provided valuable insights into race-specific resistance mediated by TKPs. We showed that an integrated HMA-like domain in WTK4 recognises AvrWTK4, and that *Bgt* can evade this recognition through point mutations or effector downregulation. These findings further highlight the versatility of HMA-like integrated domains in pathogen sensing and underscore the importance of combining genetic, molecular, and structural approaches to unravel effector-triggered immunity. From a breeding perspective, leveraging such mechanistic insights enables more informed deployment of immune receptors, and opens the door to engineering broad-spectrum resistance in cereal crops through rational design of TKPs with optimised integrated domains.

## Materials & Methods

### Fungal and plant material

All *Bgt* mutants described in this study were derived from the isolate CHE_96224 (Müller et al. 2019) and generated as previously described (Bernasconi et al. 2024). The F1 mapping population derived from the crossing of CHE_96224 and THUN-12 used for mapping the *AvrWTK4* locus was also previously described (Müller et al. 2019).

To maintain the *Bgt* isolates, the powdery mildew-susceptible wheat cultivars Kanzler (accession number K-57220) and Chancellor (accession number K-51404) were used. *Aegilops tauschii* lines (i.e. TOWWC0087, TOWWC0112, and TOWWC0154) from a diversity panel (Arora et al. 2019; Gaurav et al. 2022) containing the resistance gene *WTK4* were used to assess powdery mildew virulence on *WTK4*. For evaluating the virulence specificity of the *Bgt* mutants we used the wheat lines Axminster/8*Chancellor (*Pm1a* (Hewitt et al. 2021)), Federation*4/Ulka (*Pm2a* (Sánchez-Martín et al. 2016)), Asosan/8*Chancellor (*Pm3a*), Chul/8*Chancellor (*Pm3b*), Sonora/8*Chancellor (*Pm3c*), Kolibri (*Pm3d*), W150 (*Pm3e*), Michigan Amber/8*Chancellor (*Pm3f* (Brunner et al. 2010)), Khapli/8*Chancellor//8*Federation (*Pm4a*), Federation*8/W804 (*Pm4b* (Sánchez-Martín et al. 2021)), Kavkaz/4*Federation (*Pm8* (Hurni et al. 2013)), Amigo (1AL.1RS translocation line containing *Pm17* (Müller et al. 2022)) and the *Pm24*-containing line (kindly provided by C. Cowger).

### QTL mapping

Selected F1 progeny of the CHE_96224 x THUN-12 cross (Müller et al. 2019) were phenotyped on the *Ae. tauschii* lines TOWWC0028, TOWWC0040 and TOWWC0111 (Arora et al. 2019; Gaurav et al. 2022) and scored at 7-8 days post-infection (**Supp. Table S2**). The cultivar Kanzler was used as a wheat powdery mildew susceptible control. The phenotype was scored for individual leaf segments with a value between 0 (completely avirulent) and 100 (completely virulent), corresponding to the percentage of the leaf covered by fungal mycelium. The final score used for QTL mapping consists of an average of two to four leaf segments per *Ae. tauschii* line. QTL mapping was performed with a single interval approach using the R/qtl v.1.46.2 (https://rqtl.org/) package in R/RStudio (versions 4.3.2 and 2023.09.1, respectively). The commands used were read.cross(), jittermap() and calc.genoprobe(, step=1), as described previously (Müller et al. 2019). Single interval analysis was performed using the scanone(model=”np”) method. LOD thresholds were calculated using 1,000 permutations.

### Mutagenesis and phenotyping experiments

*Ae. tauschii* and wheat plants used for mutagenesis and phenotyping experiments were grown for 12-17 days (*Ae. tauschii*) or 9-12 days (wheat) under the following conditions: 16 h light, 8 h dark, 18°C, and 60% humidity. First leaves were then cut into 3-cm fragments and placed in Petri dishes containing water agar supplemented with benzimidazole (5 ppm, MERCK, 51-17-2). Two to four biological replicates were then spray-inoculated with *Bgt* isolates as described in our previous study (Bernasconi et al. 2024).

To identify *Bgt* mutants with gain of virulence on *WTK4*, we used the AvrXpose approach (Bernasconi et al. 2024). Briefly, the *Bgt* isolate CHE_96224 was grown on the susceptible wheat line Kanzler and irradiated with UV-C light three times. The irradiated spore mixture was then used to inoculate *Ae. tauschii* lines containing *WTK4* (TOWWC0087, TOWWC0112, and TOWWC0154). Surviving *Bgt* colonies on either one of these lines were single-spore isolated and propagated on Kanzler until sufficient spores were available for further phenotyping and DNA extraction. The phenotype of the *AvrWTK4* mutants was further evaluated on different *Pm* gene containing lines to confirm their virulence specificity to *WTK4*. Disease severity was assessed by eye 7-8 days after inoculation, and the virulence was based on the percentage of the leaf area covered by the fungal mycelium.

### DNA extraction and whole genome sequencing

The progeny of the *Bgt* F1 mapping population had already been sequenced, and a linkage map was available (Müller et al. 2019). For the newly identified *Bgt* mutants, DNA was extracted using a chloroform- and CTAB-based protocol as previously described (Bourras et al. 2015). Library preparation and Illumina whole genome sequencing were performed at Novogene (Cambridge, UK) using the NovaSeq 6000 technology, as described in our previous study (Bernasconi et al. 2024). A minimum of 12 Gb of sequence data (average coverage: 100X) was generated per *Bgt* isolate, with paired-end sequence reads of 150 base pairs (bp) and an insert size of approximately 350 bp.

### Trimming, mapping, variant calling, and transposable element detection

To prepare the Illumina reads for subsequent analysis, raw reads were first trimmed to remove Illumina oligo adapters, and sequence quality was assessed using Trimmomatic v. 0.39 (Bolger, Lohse, and Usadel 2014). Trimmed reads were then mapped to the reference genome of the *Bgt* isolate CHE_96224, genome version 3.16 (Müller et al. 2019) using bwa mem v0.7 (H. Li and Durbin 2009). We then sorted, removed duplicates, generated, and indexed the mapping (bam) files using SAMtools (H. Li et al. 2009; Danecek et al. 2021).

FreeBayes was used to perform the haplotype calling, applying the parameters -p 1 -m 30 -q 20 -z 0.03 -F 0.7 -3 200 (Garrison, Marth, and Sequencing 2012). The TASSEL5 software was used to convert the resulting vcf file into HapMap format (Bradbury et al. 2007). The software Detettore was used to detect transposable element (TE) insertions (Stritt and Roulin 2021). An in-house Python script was used to detect duplicated or deleted genes (https://gist.github.com/caldetas/24576da33d1ff91057ecabb1c5a3b6af). Variants located within 1.5 or 2 kb windows from annotated genes (for SNPs and TE insertions, respectively) were selected and visually inspected using the integrative genomics viewer software IGV (Robinson et al. 2011).

### PCR, qPCR experiments and statistical analyses

The PCR amplification of *Bgt-55150* in the *AvrWTK4* mutants was performed using the Phusion® High-Fidelity DNA Polymerase (M0530S, New England BioLabs Inc.) following the manufacturer’s recommendations, with an annealing temperature of 59 °C and an extension time of 45 seconds. The sequences of the primers used in all PCR and RT-qPCR experiments are described in **Supp. Table S6**.

RT-qPCR experiments to study the expression of *Bgt-55150* and *Bgt-646* were performed according to the MIQE guidelines (Bustin et al. 2009), following the procedure described in our previous study (Bernasconi et al. 2024). Briefly, the Dynabeads mRNA DIRECT Kit (Invitrogen) was used to extract mRNA from infected leaf material obtained from three to five independent biological replicates, two days after *Bgt* inoculation. cDNA was synthesized using the Maxima H Minus First Strand cDNA Synthesis Kit (ThermoFisher), according to the manufacturer’s instructions. The KAPA SYBR FAST qPCR kit (Kapa Biosystems) was used for the reaction. Expression was then measured in a CFX384 Touch Real-Time PCR Detection System (Bio-Rad). Gene expression was analysed using CFX Maestro software version 3.1 (Bio-Rad) and fungal glyceraldehyde 3-phosphate dehydrogenase (*GAPDH_Bgt_*) expression was used to normalize gene expression. An analysis of variance (ANOVA) followed by a Tukey HSD test was conducted to assess statistical differences between CHE_96224 and the *Bgt* mutants (**Supp. Table S7**). All analyses were performed in R using the aov and TukeyHSD functions, respectively. Finally, the glht and cld functions of the multcomp package were used to assign significance groups (Bretz, Hothorn, and Westfall 2016).

### Microscopic evaluation of immune response in *WTK4* containing *Ae. tauschii* lines

Twelve-days-old leaves of *Ae. tauschii* lines TOWWC0028 and TOWWC0077 (donors of *WTK4*) were cut into 5 cm fragments, placed on water agar Petri dishes, and infected at low density with the *Bgt* isolate CHE_96224. At 24, 48, and 96 hours post-infection (hpi), two leaves per line were destained with 3:1 ethanol:glacial acetic acid, then stained first overnight with a DAB solution (5 mM 3’3-diaminobenzidine, pH 3.8) to visualise cell death, and then for 1 minute with a Coomassie blue solution to visualise the fungal structures (0.15% Coomassie blue R-250 in methanol). The leaves were then rinsed twice with distilled water and stored in lactoglycerol (equal parts lactic acid, glycerol, and water) until microscopy. At least 100 randomly selected spores per line and time point were categorised according to the outcome of the cell attack attempt: (i) penetration attempt with consequent hypersensitive response, (ii) penetration attempt blocked by the formation of papilla, (iii) penetration and appressorium formation, and (iv) penetration and microcolony formation. Examples of these reactions can be found in **Supp. Fig. S3**. If none of these reactions occurred, or if it was not clear which reaction had occurred, the spores were assigned to the category (v), unclear reaction / no penetration.

### Cloning of expression constructs

The coding sequences of *Bgt-55150_96224_, Bgt-55150_THUN-12_, BgtE-5764_96224_* and *BgtE-5764_THUN-12_* without signal peptides (predicted by SignalP6.0; https://services.healthtech.dtu.dk/services/SignalP-6.0/) were codon-optimized for *N. benthamiana* expression with the codon optimization tool from Integrated DNA technologies (https://eu.idtdna.com) and synthesized with Gateway compatible attL sites by BioCat GmbH (https://www.biocat.com). Sequence information of all constructs produced by gene synthesis can be found in **Supp. Table S8**. The *WTK4* coding sequence was obtained from *Ae. tauschii* cDNA and can be retrieved from NCBI (GenBank: MW295405.1). Entry clones described above were cloned into the binary expression vectors pIPKb002, pIPKb004 (Himmelbach et al. 2007), and the pDEST vectors for the split luciferase assay (GW_NLUC, NLUC_GW, GW_CLUC, CLUC_GW (Gehl et al. 2011)) using the Gateway LR clonase II (Invitrogen). The constructs were subsequently transformed into *Agrobacterium tumefaciens* strain GV3101 using a freeze-thaw transformation protocol (Weigel and Glazebrook 2006).

### Cell death and interaction assays in *N. benthamiana* and wheat protoplasts

*A. tumefaciens-*mediated expression of resistance and effector genes was achieved following the procedure described previously (Bourras, Kunz, Xue, Praz, Müller, et al. 2019). For the cell death assay, *A. tumefaciens* carrying the effector or *R* gene in pIPKb004 (Himmelbach et al. 2007) were mixed in a ratio of 3:1, with the final addition of 10% volume of the P19 *A. tumefaciens* strain immediately prior to infiltration. HR development was assessed four to six days after infiltration using a Fusion FX imaging System (Vilber Lourmat, Eberhardzell, Germany). The cell death experiment was repeated three times, and a representative image was chosen for the results (**Supp. Fig. S4**). For the split luciferase assay, the effector, the *R* gene, and the P19 *A. tumefaciens* strain were mixed in equal proportions immediately prior to infiltration. Three days after infiltration, 2 leaf discs (technical replicates) from 6 biological replicates were cut, a D-luciferin substrate was added, and luminescence was measured continuously for 25 minutes at 200 ms / well using a Tecan Infinite 200 PRO LUMI plate reader.

To test cell death in wheat protoplasts a previously described protocol was used (Arndell et al. 2024). Briefly, plasmid DNA of *R* and *Avr* genes, previously cloned into pTA22 (Arndell et al. 2024), was extracted using the Xtra Midi Plus Endotoxin-free MidiPrep Kit from Macherey-Nagel. Seedlings of the *Ae. tauschii* lines TOWWC0111 and TOWWC0167, and of the wheat cultivar Fielder were grown in a growth chamber under a cycle of 12 h light (100 µmol m^-2^ s^-1^) and 12 h dark at 24 °C for 10-12 days (*Ae. tauschii*) or 7-8 days (wheat). Cell walls of epidermal peels were digested in a cellulase RS and macerozyme R-10 (Onozuka) solution, then filtered through a 40 μm nylon cell strainer and centrifuged for 3 minutes at 80 *g*. The cells were then resuspended in W5 solution (2 mM MES-KOH pH 5.7, 5 mM KCl, 125 mM CaCl_2_, 154 mM NaCl), incubated on ice, then resuspended in an appropriate volume of MMG solution (4 mM MES-KOH pH 5.7, 0.4 M mannitol, 15 mM MgCl_2_) to obtain 2-3 x 10^5^ cells per ml. Then, a DNA solution containing 3 pmol of each construct and 3 pmol of a reporter plasmid containing *YFP* was prepared. The DNA solution was then mixed in a 2 mL tube with 200 μL of the protoplast solution and ∼230 μL of the PEG solution (40% w/v PEG-4000, 0.2 M mannitol, 100 mM CaCl_2_), gently homogenised by slowly inverting the tube, and incubated for 15-30 min at room temperature. Finally, 920 μL of W5 solution was added to stop the transformation reaction. Transformed protoplasts were then centrifuged at 100 x *g* for 2 minutes, resuspended in 650 μL of W5 solution, transferred to 12-well cell culture plates, and incubated at 23°C for 16-20 hours in the dark. After incubation, 350 µL were removed, leaving the remaining protoplasts in 300 µL W5 solution. YFP fluorescence was measured in two technical replicates and three biological replicates per treatment using a plate reader. The experiment was repeated three times, for nine biological replicates in total.

### Yeast two-hybrid assay

Yeast two-hybrid assays were conducted using the MATCHMAKER GAL4 system (Clontech, USA) following the manufacturer’s instructions. The coding sequences (CDS) of full-length and truncated forms of *WTK4* were cloned into the pGADT7 vector (Clontech, USA, PT3249-5) using the *EcoRI* and *BamHI* restriction sites. The CDS of the effector genes were cloned into the pGBKT7 vector (Clontech, USA, PT3248-5) using the same restriction sites. The resulting AD and BD fusion constructs were co-transformed into *Saccharomyces cerevisiae* strain AH109 and plated on synthetic dropout (SD) medium lacking tryptophan and leucine (SD-WL) for selection. Three individual colonies from SD-WL plates were resuspended in sterile water, adjusted to an optical density at 600 nm (OD₆₀₀) of 1.0, and subsequently diluted 1:10 and 1:100. Aliquots of each dilution were plated onto SD-WL and selective SD medium lacking tryptophan, leucine, and histidine (SD-WLH), and incubated for 3-5 days at 30°C to assess protein-protein interactions.

For protein detection, two to three individual yeast colonies were inoculated into 1 mL of SD-2 liquid medium in 2 mL centrifuge tubes and incubated overnight at 30°C with shaking at 220 rpm. The cultures were centrifuged at 1000 x *g* for 5 minutes to pellet the cells. After discarding the supernatant, cell pellets were resuspended in 1 mL of 0.2 M sodium hydroxide and incubated at room temperature for 5 minutes. Cells were then pelleted again by centrifugation at 1000 x *g* for 5 minutes, the supernatant removed, and the pellets resuspended in 100 μL of protein loading buffer along with glass beads. The samples were vortexed for 5 minutes to lyse the cells, followed by heating at 95°C for 10 minutes. The resulting protein extracts were subsequently used for immunoblotting analysis.

### Protein detection for split-luciferase assay

The soluble protein fraction was extracted from *N. benthamiana* leaves 3 days after infiltration with *A. tumefaciens*. Six leaf discs in total (0.5 mm diameter) from 3 infiltrated leaves were pooled, ground in liquid nitrogen, and resuspended in 2x modified Laemmli-Buffer (100 mM Tris-HCl pH 6.8, 200 mM DTT, 0.04% bromophenol blue, 20% glycerol, 2% SDS), boiled at 90°C for 5 minutes and centrifuged at 13,000 rpm for 1 minute. Twenty microlitres of total protein extract were separated in SDS polyacrylamide (PA) gels (7.5% or 12%) and semi-dry blotted to a Nitrocellulose membrane using a Trans-Blot SD Semi-Dry Transfer Cell (Bio-Rad). For LUC tag detection, the primary antibody (anti-LUC, rabbit polyclonal, Sigma-Aldrich) was used at a dilution of 1:3,000, membranes washed with TBST and then incubated with anti-rabbit peroxidase antibody (anti-rabbit-HRP, mouse, LabForce) diluted 1:3,000. Peroxidase chemiluminescence was detected using a Fusion FX Imaging System (Vilber Lourmat, Eberhardzell, Germany) after application of a WesternBright ECL HRP substrate (Advansta). The membranes were also incubated with Ponceau stain for total protein detection.

### Structural modelling and other bioinformatic analyses of HMA_WTK4_

The 3D modelling of AvrWTK4 and WTK4 was performed using AlphaFold2 and AlphaFold3 (Jumper et al. 2021; Mirdita et al. 2022), and the interaction between AvrWTK4 and WTK4 using AlphaFold multimer through ColabFold (Mirdita et al. 2022) or the AlphaFold server (https://alphafoldserver.com/). The protein sequences from tandem kinase proteins RPG1, YR15, SR60, and SR62^TK^ have been retrieved from UniProt (https://www.uniprot.org/). HIPP39 was found through blastp of HMA_WTK4_ with standard parameters in the NCBI database (https://blast.ncbi.nlm.nih.gov/). Their 3D structures were predicted with AlphaFold2 and visualised using PyMOL. The HMA-like domain of WTK4 was annotated following the ProSite automated annotation, and the HMA-like domain of RPG1 was annotated using the multiple sequence alignment of the HMA domains of known R proteins and HMA_WTK4_, shown in **Supp. Fig. S4** (performed using the Clustal Omega software; https://www.ebi.ac.uk/jdispatcher/msa/clustalo). TM-align was used to infer structural similarity between HMA_WTK4_ and RPG1, RGA5, Pikp1 and HIPP39, using their pdb structure files as input (Zhang 2005). Any structure with a TM score greater than 0.6, indicating a 60% structural similarity, was considered a structural homologue.

## Acknowledgements

We thank Gerhard Herren for the help with RT-qPCR and for technical support, Lukas Kunz for fruitful discussions and material, Karl Huwiler for greenhouse support, Megan Outram, Melania Figueroa and Peter Dodds for the support in establishing the protoplast cell death assay.

## Author contributions

Z.B., J.S.-M. and B.K., conceived the study and the experiments, interpreted results, and wrote the manuscript, with inputs from T.W. and B.B.H.W. Z.B. and U.S. identified the *Bgt* mutants, performed the phenotyping experiments, performed and interpreted the RT-qPCR experiments. A.H. and Z.B. performed the protoplast cell death assay. Z.B., Y.X., R.C. and V.W. performed the protein-protein interaction assays. M.L. performed the microscopy analysis of infected *Ae. tauschii* leaves. M.H., T.W. and M.C.M. contributed to the bioinformatic analyses. All authors have read and agreed to the published version of the article.

## Data availability

Raw FASTA sequences of all *Bgt* mutants generated in this study are available through NCBI (BioProject ID: PRJNA1016363).

