## Supplementary Figures for "An HMA-like integrated domain in the wheat tandem kinase WTK4 recognises an RNase-like pathogen effector"

| Isolate | CHE_96224 | AvrWTK4_2 | AvrWTK4_4 | AvrWTK4_5 | AvrWTK4_6 |
|---|---|---|---|---|---|
| *Bgt-646* | intact | **mutated** | intact | intact | intact |
| *Bgt-55150* | intact | intact | **mutated** | intact | **mutated** |

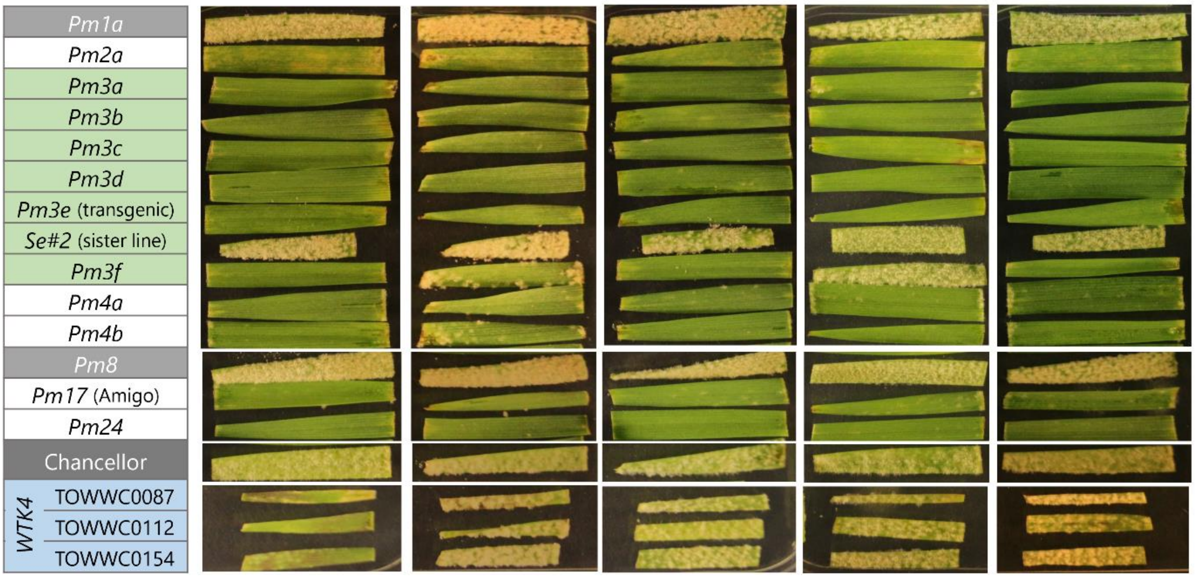


**Supp. Fig. S1**. ***AvrWTK4* mutants are virulent on *WTK4*-containing lines**. The *AvrWTK4* mutants exhibit specific gain of virulence on three different *Ae. tauschii* lines (TOWWC0087, TOWWC0112, and TOWWC0154), but no virulence on near-isogenic lines (NILs) carrying various *R* genes for which their parental isolate CHE_96224 is avirulent. The exception is mutant AvrWTK4_5, which exhibits additional gain of virulence on *Pm3f*, caused by an unknown mutation. Cultivars susceptible to CHE_96224 (the *Pm1a* NIL, the sister line of the transgenic *Pm3e* line (here called *Se#2*), the *Pm8* NIL and Chancellor) are highlighted in grey. NILs containing *Pm3* alleles are depicted in green, and the three *Ae. tauschii* lines containing *WTK4* are in blue. The remaining *Pm* lines are shown in white.

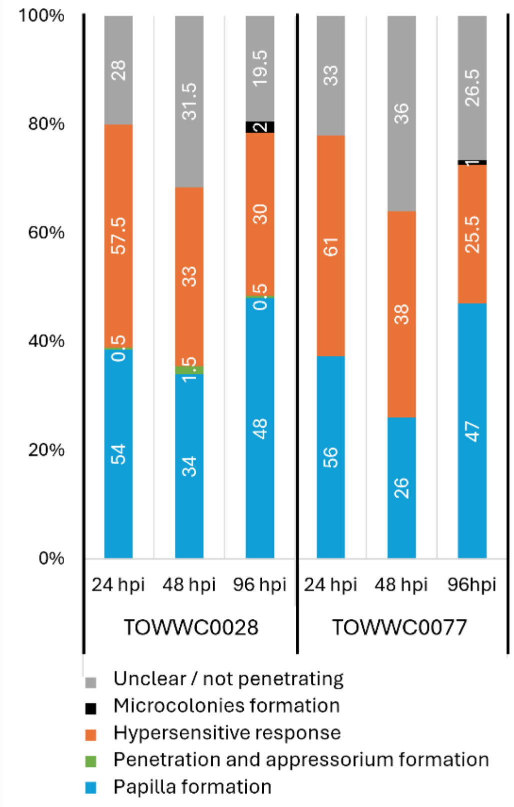


**Supp. Fig. S2. Microscopic evaluation of the immune response induced by the avirulent *Bgt* isolate CHE_96224 on two *WTK4*-containing *Ae. tauschii* lines (TOWWC0028 and TOWWC0077).** At least 100 spores per leaf, time point, and line were visually inspected. Coomassie staining was used to visualise fungal structures (blue/purple), and DAB staining was used to identify cells undergoing hypersensitive response (HR) (dark brown). The number of *Bgt* spores causing HR (orange in the graph), blocked by papilla formation (blue), penetrating and creating an appressorium (green) or forming a microcolony (black) were counted. If the reaction was unclear, spores were classified as “Unclear / not penetrating”. Examples of a *Bgt* spore attempting to penetrate a plant cell undergoing HR and of a *Bgt* spore attempting to penetrate a plant cell blocked by papilla formation are shown in **Fig. 3A-B**, respectively.

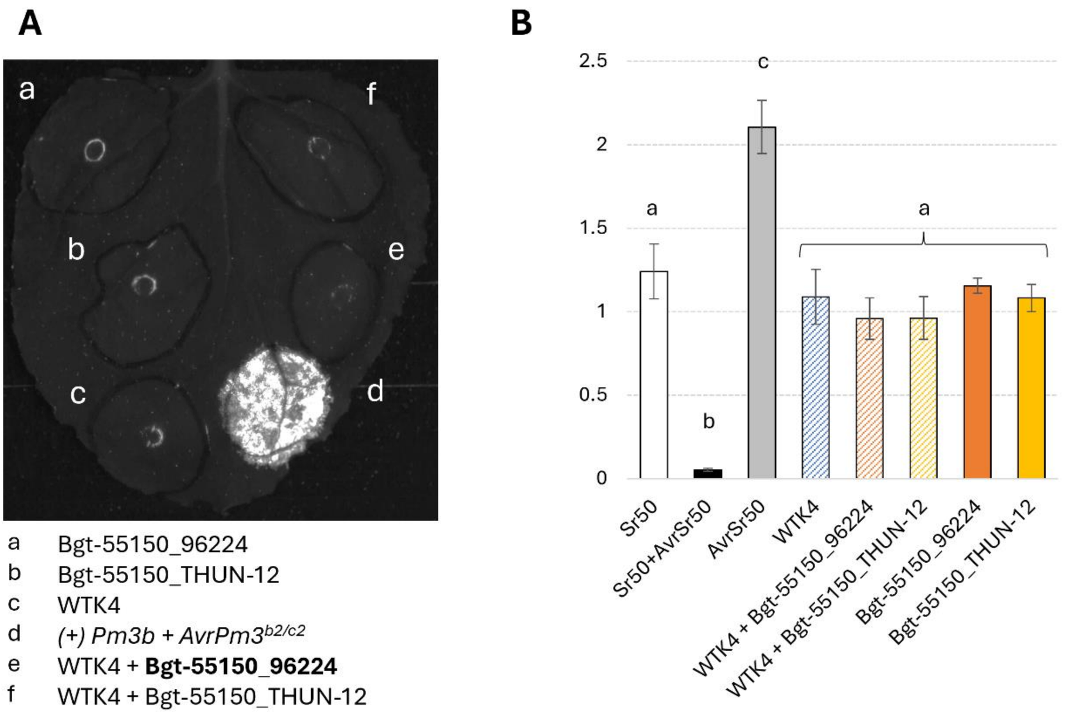


**Supp. Fig. S3. Co-expression of *AvrWTK4$_{96224}$* (*Bgt-55150_96224*) and *WTK4* does not lead to cell death in (A) *N. benthamiana* or (B) wheat protoplasts (wheat cv. Fielder).** The cell death signal in *N. benthamiana* was measured 5 days post-infiltration. The YFP fluorescence in wheat protoplasts was measured 20 hours post-transfection and normalised to the fluorescence value of protoplast transfected with YFP alone. As a positive control, *Sr50* co-transfected with its corresponding *Avr* effector *AvrSr50* was used. Two technical and three biological replicates were measured per transfection. Significance and p values (significance threshold of $p < 0.05$) were calculated using a two-way analysis of variance (ANOVA) followed by a Tukey's HSD test (**Supp. Table S7**). Bars with the same letter are not significantly different.

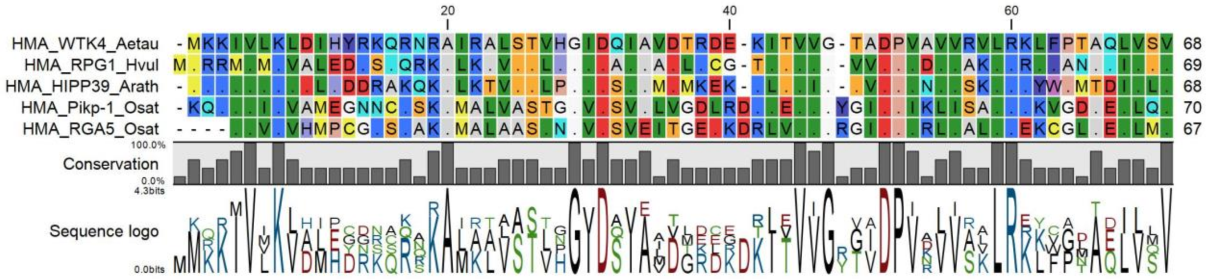


**Supp. Fig. S4**. **Protein alignment of the HMA domains of WTK4 (from *Ae. tauschii*), RPG1 (from barley), Pikp-1, RGA5 (both from rice), and HIPP39 (the closest $HMA_{WTK4}$ domain homologue from *Arabidopsis thaliana*)**. The amino acid colouring reflects traditional biochemical properties of the amino acids (e.g. polarity) and shows a high degree of conservation between the different HMA domains. The HMA domain sequences were extracted from the UniProt database or by blasting the HMA domain of WTK4 in the NCBI database and aligned using Clustal Omega.

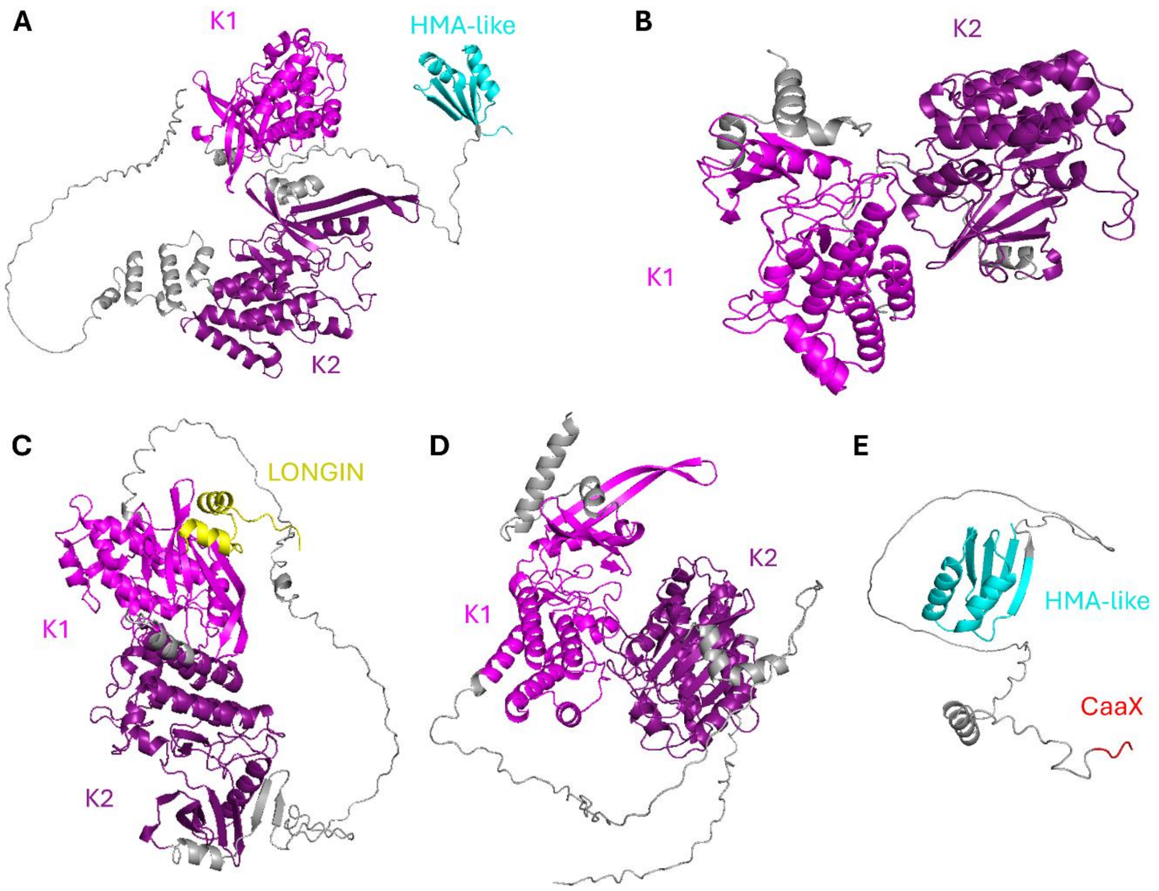


**Supp. Fig. S5**. **Protein structure prediction of TKPs and the HMA-domain containing protein HIPP39.** (**A**) RPG1, (**B**) Yr15 (WTK1), (**C**) Sr60 (WTK2), (**D**) $Sr62^{TK}$ (WTK5) and (**E**) HIPP39. The 3D structure prediction for other tandem kinase proteins, i.e. WTK6-vWA, WTK7-TM and RWT4 (allelic to Pm24) can be found in the studies of Wang et al. (2023), Li et al. (2024) and Sung et al. (2025), respectively. The HMA-like domains are shown in cyan and consist of the typical βαββαβ 3D structure. The two kinase domains of the tandem-kinase proteins (TKPs) are indicated as K1 (light magenta) and K2 (dark magenta) based on their order (from the N-terminus) in the protein. K1 is a pseudokinase in WTK4 and RPG1, whereas it is an active kinase in all other TKPs. K2 is an active kinase for WTK4, RPG1 and Sr60 (which has two active kinases), and a pseudokinase for the other WTKs. In addition, Sr60 has a predicted N-terminal LONGIN domain, possibly involved in association with vesicular membrane proteins (in yellow, predicted with low confidence by ScanProsite). HIPP39 has a C-terminal isoprenylation motif (CaaX, in red; A being an aliphatic amino acid, and X any amino acid). The linker sequences (without any domain prediction) between the different domains are shown in grey. All structures have been predicted with AlphaFold2 and visualised with PyMOL.

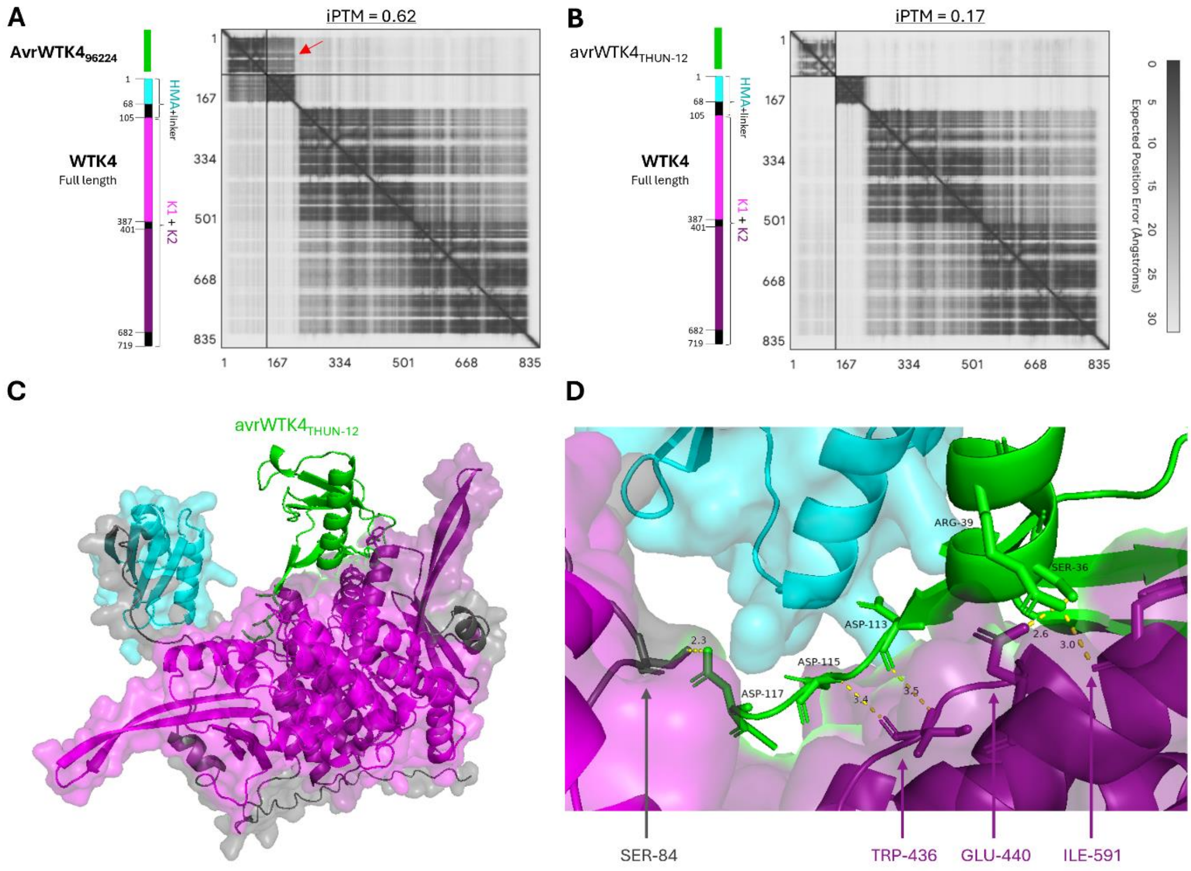


**Supp. Fig. S6. The prediction of the WTK4 interaction with avrWTK4$_{THUN-12}$ differs from the interaction with the avirulent effector variant AvrWTK4$_{96224}$.** Predicted alignment errors (PAE) of (**A**) AvrWTK4$_{96224}$-WTK4 (ipTM value 0.62) and of (**B**) avrWTK4$_{THUN-12}$-WTK4 (ipTM value 0.17) by AlphaFold3. The region of interaction (exhibiting low PAE, dark grey) for AvrWTK4$_{96224}$-WTK4 is indicated with a red arrow and corresponds to HMA$_{WTK4}$. For avrWTK4-WT4, the PAE is high along all the interaction surfaces, indicating low confidence of interaction. Panels (**C**) and (**D**) show that AlphaFold3 predicts the avrWTK4$_{THUN-12}$-WTK4 interaction to occur through a different surface compared to AvrWTK4$_{96224}$-WTK4 interaction (**Fig. 4**). The interaction interface between WTK4 and avrWTK4$_{THUN-12}$ is predicted to involve the K1 and K2 domains of WTK4 and the effector residues S36, R39, A113, A115, and A117 (with low confidence, see panel **B**). Effector residues 30 and 33, which interacted with HMA$_{WTK4}$ in AvrWTK4$_{96224}$, are not involved in the interaction in avrWTK4$_{THUN-12}$. The interacting residues of WTK4 are shown below the image, while the interacting residues of avrWTK4$_{THUN-12}$ and the distance in ångström are shown within the image.

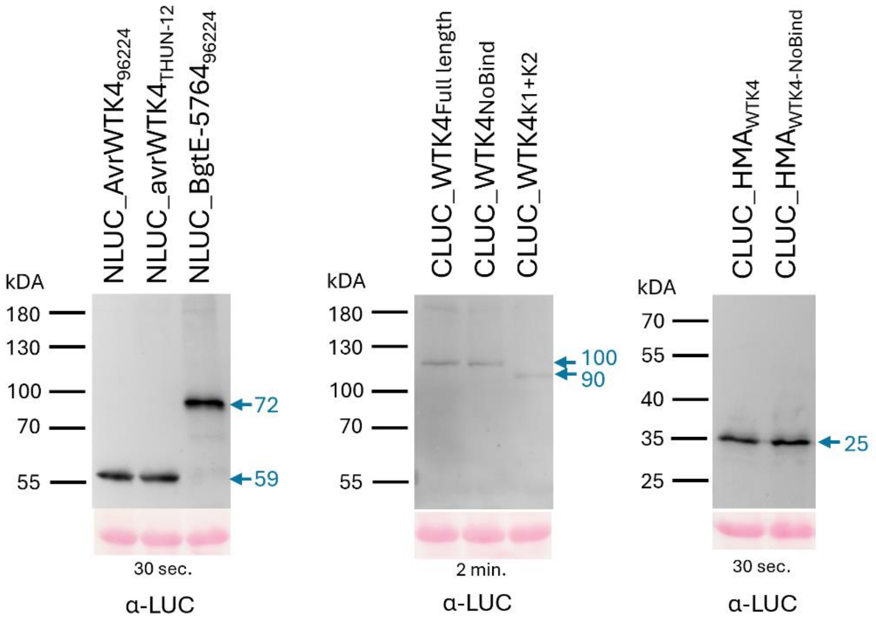


**Supp. Fig. S7. Protein detection of the proteins used in the split-luciferase experiment**. LUC-tagged constructs used in the split-luciferase assay were infiltrated into *Nicotiana benthamiana* leaves along with P19, using a construct-to-P19 ratio of 2:1. Three days after infiltration, six leaf discs were collected from three different leaves, and total protein was extracted as outlined in the Methods section. The WTK4 full length and K1+K2 constructs were exposed for 2 minutes, while the Avr and HMA+linker constructs for 30 seconds. The expected sizes in kDa are depicted in blue on the right of every blot. No significant expression difference was detected between WTK4 and WTK4$_{NoBind}$, HMA$_{WTK4}$ and HMA$_{WTK4-NoBind}$, and AvrWTK4$_{96224}$ and avrWTK4$_{THUN-12}$. Below, membranes stained with a Ponceau solution as a loading control.

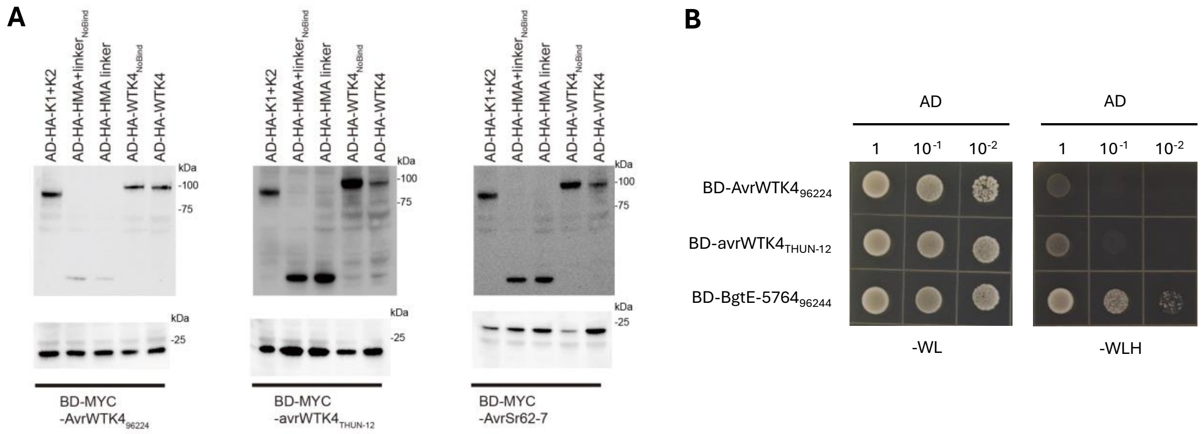


**Supp. Fig. S8**. **Protein detection of yeast two-hybrid constructs and autoactivity of BgtE-5764.** (**A**) Immunoblot detection of protein expressed in yeast in **Fig. 4E**, detected using anti-HA or anti-Myc antibodies. The coding sequences of the HA and Myc tags are integrated into the AD and BD vectors, respectively, and are in the same reading frame as their corresponding fusion proteins. (**B**) The BD-BgtE-5764$_{96224}$ fusion protein exhibited autoactivation activity in yeast. BD-BgtE-5764$_{96224}$ was co-transformed with the empty vector pGADT7 into the yeast strain AH109, it grew normally on the selective medium SD-3, indicating that the BgtE-5764$_{96224}$ fusion protein is unsuitable for use as a negative control in yeast two-hybrid experiments.
